# ANCHOR, a technical approach to monitor single-copy locus localization *in planta*

**DOI:** 10.1101/2021.03.08.434382

**Authors:** Anis Meschichi, Mathieu Ingouff, Claire Picart, Marie Mirouze, Sophie Desset, Franck Gallardo, Kerstin Bystricky, Nathalie Picault, Stefanie Rosa, Frédéric Pontvianne

**Affiliations:** Swedish University of Agricultural Sciences, Uppsala, Sweden; Institut de Recherche pour le Développement, Université de Montpellier, UMR232, 34394 Montpellier, France; CNRS, Laboratoire Génome et Développement des Plantes (LGDP), Université de Perpignan Via Domitia, Perpignan, France; CNRS, GReD, Université Clermont Auvergne, INSERM, BP 38, 63001 Clermont–Ferrand, France; NeoVirTech SAS, 1 Place Pierre Potier, 31000 Toulouse, France; Laboratoire de Biologie Moléculaire Eucaryote (LBME), Centre de Biologie Intégrative (CBI), CNRS, UPS, University of Toulouse, 31062, Toulouse, France

**Author notes:** equal contribution (Anis Meschichi, Mathieu Ingouff).

## Abstract

Gene expression is governed by several layers of regulation which in addition to genome organization, local chromatin structure, gene accessibility and the presence of transcription factors also includes gene positioning. Although basic mechanisms are expected to be conserved in Eukaryotes, surprisingly little information on the role of gene positioning is available in plant cells, mainly due to the lack of a highly resolutive approach. In this manuscript, we adapted the use of the ANCHOR system to perform real-time single-locus detection *in planta.* ANCHOR is a DNA-labelling tool derived from the partitioning system. We demonstrate its suitability to monitor a single-locus *in planta* and used this approach to track chromatin mobility during cell differentiation in Arabidopsis root epidermal cells. Finally, we discuss the potential of this approach to investigate the role of gene positioning during transcription and DNA repair in plants.

## INTRODUCTION

In Eukaryotes, genetic information is encoded in the chromatin, a complex structure composed of DNA packed around an octamer of histones in the nucleus. Chromosome territories form large compartments in the nuclear interior, themselves containing chromatin domains harbouring different epigenetic signatures (Santos et al., 2020; Pontvianne and Grob, 2020; Nguyen and Bosco, 2015). In these domains, gene positioning and accessibility is very dynamic in response to several key biological processes that include gene transcription, genome replication and DNA repair for example. Fluorescence *in situ* Hybridization (FISH) approaches such as padlock-FISH enable to detect a single-copy locus using fixed material (Feng et al., 2014). However, imaging techniques using non-living organisms is insufficient to track spatial and temporal dynamics of loci. Live-cell imaging approaches enable to visualize gene positioning during these various processes, bringing key elements for their understanding (Shaban and Seeber, 2020; Dumur et al., 2019).

Microscopic detection of genomic loci in plants is possible through the use of different strategies, including zinc-finger, TALE and CRISPR/Cas9 based imaging (Lindout et al., 2007; Fujimoto et al., 2016; Khosravi et al., 2020). Unfortunately, these techniques have been restricted to follow the dynamics of highly repeated regions (centromeric repeats, telomeric sequences and ribosomal RNA genes). Monitoring of a single locus in living plants is possible thanks to the addition of *lacO* motifs to which the transcription factor LacI, fused to a fluorescent protein, can bind (Fang and Spector, 2007; Kato and Lam, 2003). Live-cell imaging of *FLOWERING LOCUS C (FLC)* alleles associated to *lacO (FLC-LacO)* could be performed to demonstrate that *FLC-LacO* repression during vernalization provokes their physical clustering (Rosa et al., 2013). Moreover, the repressor protein Tet fused to a fluorescent protein could also be used to labelled a genomic region containing numerous Tet operator sequences (Matzke et al., 2005). In both cases, amplification of the signal is directly linked to the multiplicity of the targeted sequences. However, these repetitions often affect local chromatin organization, leading to local bias (Watanabe et al., 2005). Thus, a standardized and robust technique to track single locus dynamics is still not yet available.

The ANCHOR system is a DNA-labelling tool derived and optimized from bacterial partition complexes. A single-copy of *parS* −1 kb long fragment-serves as a binding platform for ParB proteins binding (Dubarry et al., 2006). Natural *ParS* sequence is composed of 4 canonical inverted repeat sequences that are bound via the helix-turn-helix (HTH) motif present in ParB (Funnell, 2016). Upon binding, oligomerization of ParB proteins then spreads onto the ParS sequence and adjacent DNA (Figure 1A). Importantly, oligomerized fluorescent ParB are loosely associated and can be displaced transiently and easily upon transcription or DNA repair (Saad et al., 2014). This phenomenon is also described as a caging step (Funnell, 2016). This system has been used successfully to visualize a unique locus in living yeast and human cells (Germier et al., 2017). This approach is also largely used to monitor DNA virus positioning in human cells (Gallardo et al., 2020; Mariamé et al., 2018; Komatsu et al., 2018; Blanco-Rodriguez et al., 2020; Hinsberger et al., 2020). In this manuscript, we demonstrate that the ANCHOR system can also be adapted to visualize a single-locus in fixed and living plant tissues. Using this approach, we also show that chromatin mobility is different in differentiated versus meristematic cells *in planta.*

**Figure 1:**
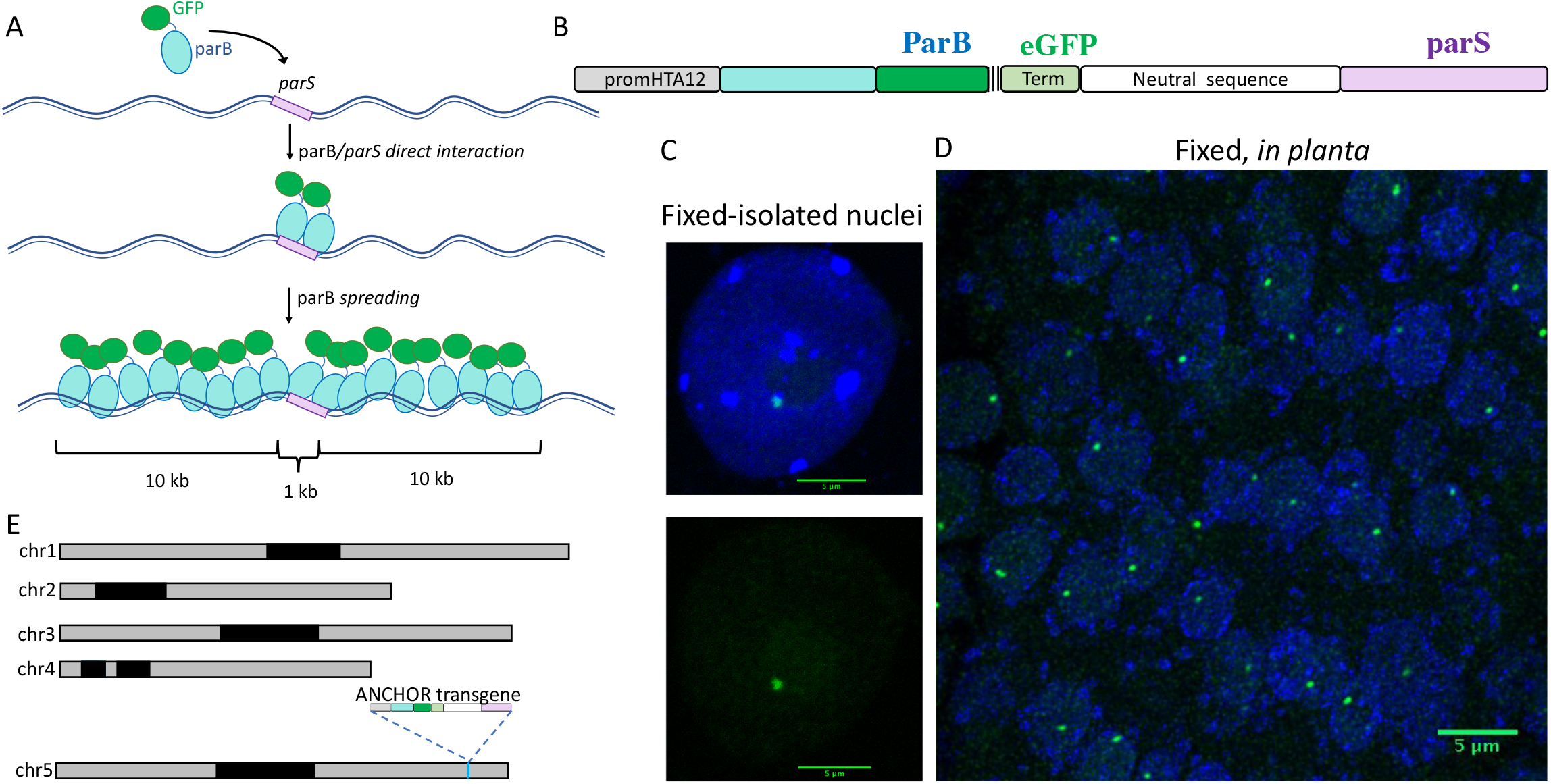
Establishment of a protocol allowing the use of the ANCHOR system i*n planta*. A. Schematic representation of the ANCHOR system. ParB proteins fused to GFP can directly bound to *parS* sequence as a dimer. *parS*-ParB interactions provoke a conformational change in ParB proteins that induce their oligomerization along the flanking genomic region. B. Cassette used to transform *Arabidopsis thaliana* Col-0 plants to test the ANCHOR system *in planta.* A strong promoter is used to expressed the ParB protein fused to an enhanced GFP and three FLAG tags. After a Terminator sequence, a 1500 pb long spacer sequence has been added to separate the ParB reporter line and the 1 kb-long *parS* sequence. Detection of a *parS*-ParB:GFP focus (Green) in an isolated leaf nucleus (C) and in fixed root tissues (D) of *A. thaliana* plants containing the ANCHOR cassette described in B. Nuclear DNA is labelled with DAPI (blue). Bar = 5 μm. E. Position of the transgene in the ANCHOR line T2F in the Arabidopsis genome using Nanopore sequencing. The transgene presented in B is inserted on chromosome 5, position 23,675,998 pb.

## MATERIAL AND METHODS

### Plant Materials and Growth Conditions

*Arabidopsis thaliana* ecotype Col-0 was used in this study. *lacO*/LacI line used comes from the following source (Matzke et al., 2005).

Plants were transformed by agroinfiltration using the floral dip protocol (Clough and Bent, 1998), using *Agrobacterium tumefaciens* GV3101 strain. Transformants were selected on Basta (10 mg/L). All plant material used here was grown in control growth chambers on soil at 21°C with a daylight period of 16 hr/day. T2F line was crossed to Col-0 plant expressing the Histone variant H2AW fused to RFP (Yelagandula et al., 2014).

For MSD experiments seeds were surface sterilized in 5% v/v sodium hypochlorite for 5 min and rinsed three times in sterile distilled water. Seeds were stratified at 4°C for 48 h in the darkness and plated on Murashige and Skoog (MS) medium. Seedlings were placed in a growth cabinet (16 hours light, 22°C) for 1 week in vertically oriented Petri dishes before imaging.

### Plasmid construction

A cassette allowing the expression of ParB has been synthetized by Genescript^®^. The nature and sequences of the ANCHOR system are confidential and the property of NeoVirTech SAS. The cassette was cloned into the pEarleyGate302 vector (Earley et al., 2006).

### Cytogenetic Analyses

DAPI-stained nucleus analyses were performed using nuclei from leaves of 3- or 4-week-old plants as described previously (Pontvianne et al., 2012). Nuclei with different ploidy levels were isolated with a S3 cell sorter (Biorad^®^), as described in (Pontvianne et al., 2016), except that propidium iodide was used to stain nuclei. Immunolocalization experiments were performed as described previously (Durut et al., 2014) using anti-H3K27me3 or anti-H3Ac antibodies (Abcam) to a 1/1000 dilution. Zeiss LSM 700 confocal was used to generate Figure 1, while Zeiss LSM 800 with an Airyscan module was used to generate images from Figure 2 and 4 with a 63x objective, N.A. 1.4 and pixel size 0.028×0.028×0.160 μm^3^. Live-cell imaging presented in Figure 3 were performed using a spinning disk Zeiss Cell observer equipped with a high-speed Yokogawa CSUX1spinning disk confocal, an ORCA-flash 4.0 digital camera (Hamamatsu) and a ×40 water objective N.A. 1.2. GFP was excited at 488 nm.

**Figure 2:**
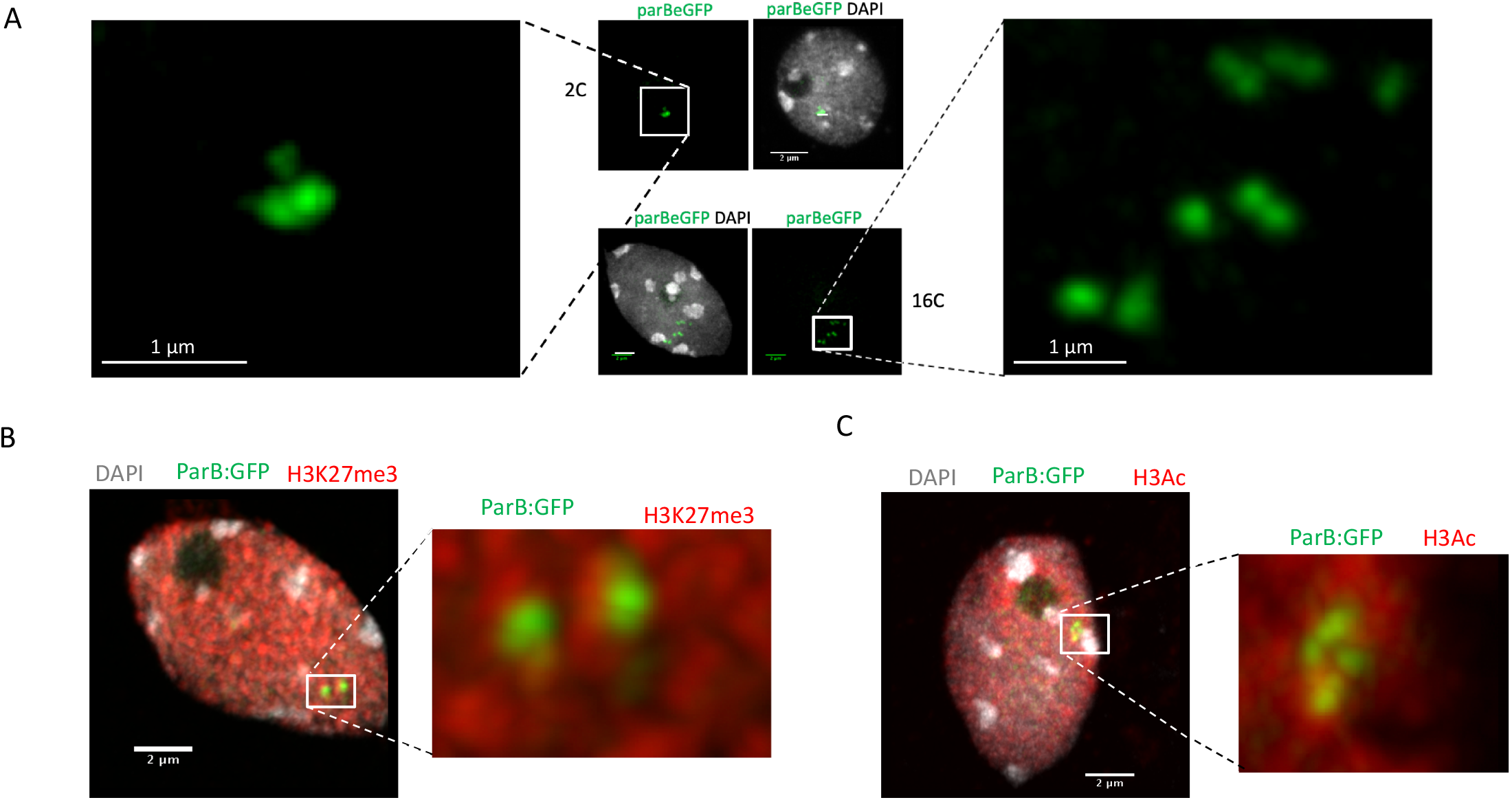
ANCHOR system is suitable for multiple allele detection and can be coupled with immunolocalization analyzes. A. Detection of *parS*-ParB:GFP foci (Green) in fixed and sorted nuclei according to their ploidy levels by Fluorescent-Assisted Cell Sorting (FACS). Nuclear DNA is labeled with DAPI (grey). Enlarged view of the *parS*-ParB:GFP foci are presented to facilitate signal visualization. Bar = 1 μm. B-C. Detection of *parS*-ParB:GFP foci (Green) and post-transitionally modified histones (red) in fixed and isolated nuclei *from A. thaliana* Col-0 plants T2F. Nuclear DNA is labeled with DAPI (grey). Trimethylated H3K27 signals are shown in the panel B, while acetylated H3 are shown in the panel C. Enlarged views of the *parS*-ParB:GFP foci are presented to facilitate signal visualization. Bar = 2 μm.

### Nanopore sequencing

Genomic DNA preparation was performed as previously described (Picart-Picolo et al., 2020). Library preparation was performed using the 1D Genomic DNA by ligation kit SQK-LSK109 (Oxford Nanopore Technologies), following the manufacturer’s instructions. The R9.5 ONT flow-cell FLO-MIN106D (Oxford Nanopore Technologies) was used. We obtained 1.93 Gb of sequences (11X coverage) with an average read length of 3, 675kb for ANCHOR T2F line. ONT reads mapping the transgene were mapped, filtered and aligned using Geneious software (Kearse et al., 2012).

### Live cell Imaging

For figure 5, time-lapse imaging of Arabidopsis roots has been carried out using a Zeiss LSM 780 confocal microscope using a 63x water immersion objectives (1.20 NA). For visualization of root cell contours stained with propidium iodide, an excitation line of 488 nm was used and signal was detected at wavelengths of 580 to 700nm. For observation of GFP expression, we used respectively a 488-nm excitation line and a BP filter of 505-550 nm. For all experiments, images were acquired every 6 s taking a series of 3 optical sections with Z-step of 2 μm for 5 min. Each movie has a format of 512 × 512 pixels and a 3× zoom factor.

The 7-d-old seedlings were mounted in water, or propidium iodide, between slide and cover slip and sealed with 0.12-mm-thick SecureSeal Adhesive tape (Grace Bio-Labs), to avoid root movements and drying during imaging.

### Mean square displacement analysis

All the movies have been analyzed with Fiji software (NIH, Bethesda, MD, http://rsb.info.nih.gov/ij/) and with the plugin SpotTracker 2D (obtained from http://big-www.epfl.ch/sage/soft/spottracker). MSD analysis was performed as described in Meschichi et al. 2021. All quantitative measurements represent averages from at least 9 cells. From the MSD plot, we calculated the radius of constraint by the square root of the plateau of the MSD curve multiplied by 5/4. The standard Mann-Whitney U-test test was used to determine the statistical significance of results. For statistical analysis, we used the GraphPad Prism 8.3 software.

## RESULTS

### Development of the ANCHOR system

Our goal was to adapt and facilitate the use of the ANCHOR system in plants. We therefore combined in a single transgene the two elements of the ANCHOR system (ParB and its target sequence *parS*). A codon optimized *ParB* gene for Arabidopsis was fused to an in frame enhanced Green Fluorescent Protein (GFP) and triple FLAG tag sequence to allow detection in living and fixed nuclei (Figure 1B). This expression cassette was placed under the control of a housekeeping promoter. At the 3’ end of the ParB construct we added the 1kb long ParB target sequence *parS* separated by a 1,5 kb long spacer sequence to prevent potential interference of *ParB* gene transcriptional activity. Such design allows a rapid selection of transgenic plants containing both linked ANCHOR elements without the need to combine them and to prevent their segregation when crossed to other genetic background. Second, if we can detect a signal, then it would suggest that the transcriptional activity leading to the expression of ParB:eGFP is not perturbed by the local caging of ParB:GFP proteins.

Wild-type Col-0 plants were transformed with the transgene and selected using Basta herbicide. Isolated fixed nuclei from leaves eight different T1 transformants revealed the presence of *parS*-ParB:GFP foci in five of them (Figure 1C). To test the robustness of the detection approach, we then analysed the entire root-tip from one ANCHOR line comprising a single copy insertion at generation T2 (Figure 1D). One *parS*-pParB:GFP signal was detectable in almost all nuclei analysed. Importantly, signal to noise ratio was quite high (Figure 1D).

To further characterize the ability of the ANCHOR system to follow a single-locus *in planta,* it is important to know the exact location of the transgene. We performed long-read Nanopore sequencing on one ANCHOR line with one single insertion (T2F), and extracted all long reads corresponding to the transgene to map its location in the genome. Sequence analyses revealed that the transgene could be located on the lower arm of chromosome 5, at position 23 675 998, in an intergenic region (Figure 1E). This position is flanked by a region enriched in active chromatin marks and a region enriched with Histone 3 trimethylated Lysine 27 (H3K27me3), a repressive mark deposit by the Polycomb repressive complex 2 (PRC2) (Figure S1) (Sequeira-Mendes et al., 2014).

### Detection of *parS*-ParB *foci* in fixed cells

In root tip cells, as presented in (Figure 1D), only 2C diploid cells can be found (Hayashi et al., 2013). One unique spot was usually detected in those cells, sometimes appearing as a doublet. Because the ANCHOR system is based on protein aggregation, we wondered whether analysing ANCHOR signals in endoreplicated cells would lead to an increase number of detected foci. We isolated 2C and 16C cells by fluorescent-assisted cell sorting using propidium iodide labelling. We stained sorted nuclei with DAPI and counted *parS*-ParB:GFP signals per nuclei. We found that the number of detected signals is higher in sorted nuclei presenting a higher endoreplication rate (Figure 2A). These data suggest that the ANCHOR system is suitable to detect multiple loci simultaneously, and that this reporting system does not lead to aberrant locus aggregation.

In the T2F line, the transgene is located on an arm of the chromosome 5, in a region enriched in H3K27me3 deposited by the PRC2, but flanked by a genomic region enriched by active chromatin marks (Figure S1). We therefore took advantage of this peculiar chromatin environment to test the possibility to combine both immunostaining and *parS*-ParB:GFP signals detection. Immunostaining experiments were performed on isolated leaf nuclei from 3 week-old plants using either an antibody against Histone 3 acetylated (H3Ac) active mark or H3K27me3. As expected the tested histone marks and *parS*-ParB:GFP signals are excluded from heterochromatic foci easily visible by DAPI staining and corresponding to the centromeric, pericentromeric and nucleolus organizer regions (Figure 2B-C). Although no clear overlap could be detected between *parS*-ParB:GFP signals and H3K27me3 marks, at least partial overlap can be seen between *parS*-ParB:GFP signals and H3Ac marks (Figure 2B-C and S3). This result is expected since active transcription is necessary to produce ParB:GFP proteins.

### Detection of *pαrS*-ParB *foci* in live-cell imaging

Previous studies demonstrated that global genome organization can be cell specific and vary depending on the developmental stage analysed (Pontvianne and Liu, 2019). We therefore tested our ability to detect *parS*-ParB:GFP signal in different cell-types, directly *in planta*. Because autofluorescence can be high in certain plant cells, we crossed our T2F line with another *A. thaliana* Col-0 line expressing the Histone 2A variant H2AW, fused to the Red Fluorescent Protein (RFP) to allow heterochromatin visualization directly in living cells (Yelagandula et al., 2014). Plants were grown on MS media directly in petri dish compatible with confocal imaging. We analysed several tissues, including meristematic and differentiated root cells, leaf cells, trichome cells and pollen grains. For tested materials, we were always able to visualize *parS*-ParB:GFP signals (Figure 3 and S2). As expected, *parS*-ParB:GFP signals are excluded from heterochromatin area. Note that the nuclear area can sometimes be seen due to non-associated ParB proteins that are diffusing in the nucleoplasm.

**Figure 3:**
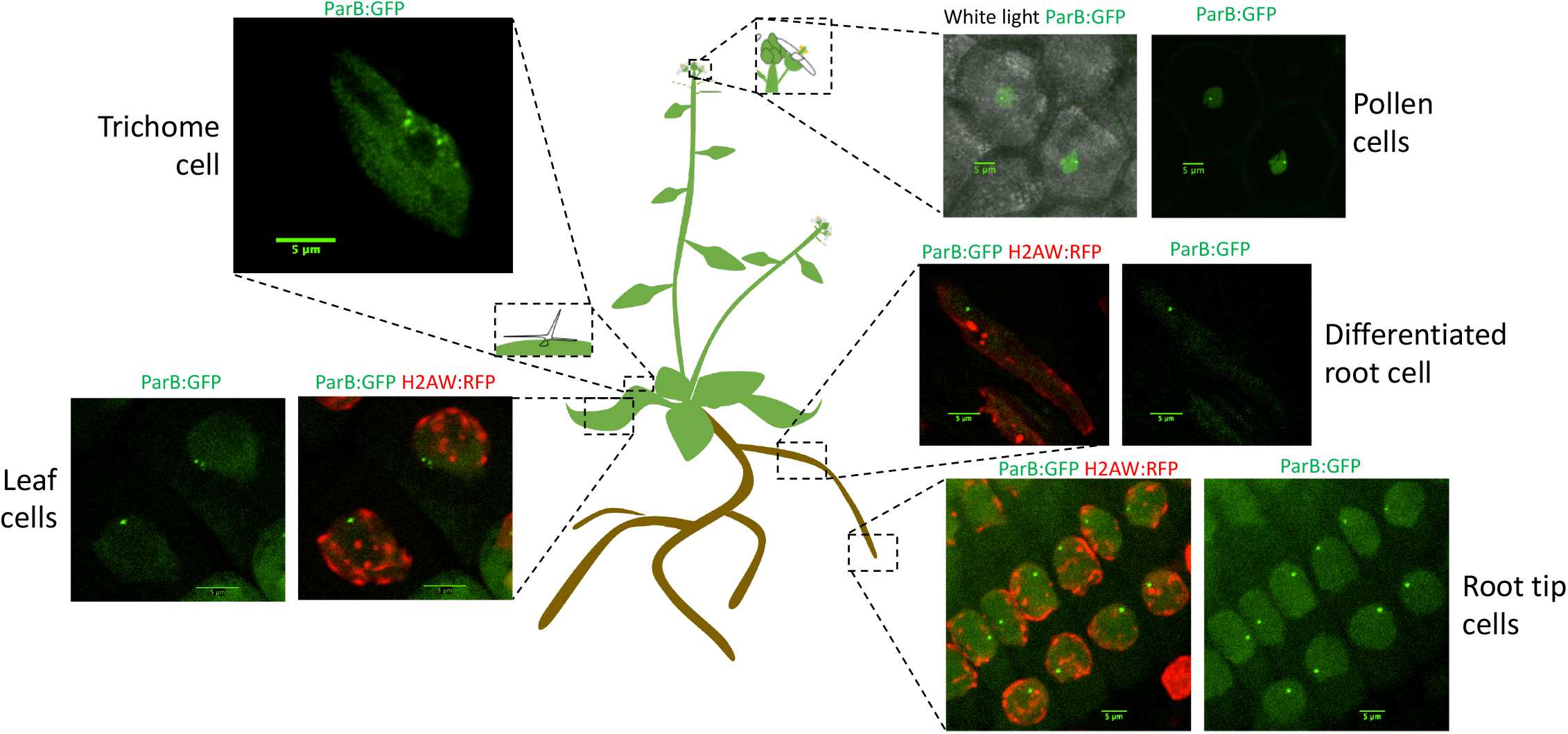
ANCHOR system is suitable to monitor a single-copy locus in live and in different tissues. Schematic representation of an *Arabidopsis thaliana* plant to illustrate the different tissues tested to monitor parS-ParB:GFP signals by live-cell imaging. ParB:GFP signals are in green and Histone variant H2AW:RFP is shown in red.

The ANCHOR system does not require high DNA accessibility to allow *parS*-ParB:GFP signals visualization. In a highly condensed chromatin context like during mitosis, we could still to detect *parS*-ParB:GFP signals in condensed chromosomes, although signal is usually less bright than in the neighboring cells (Figure 4A).

**Figure 4:**
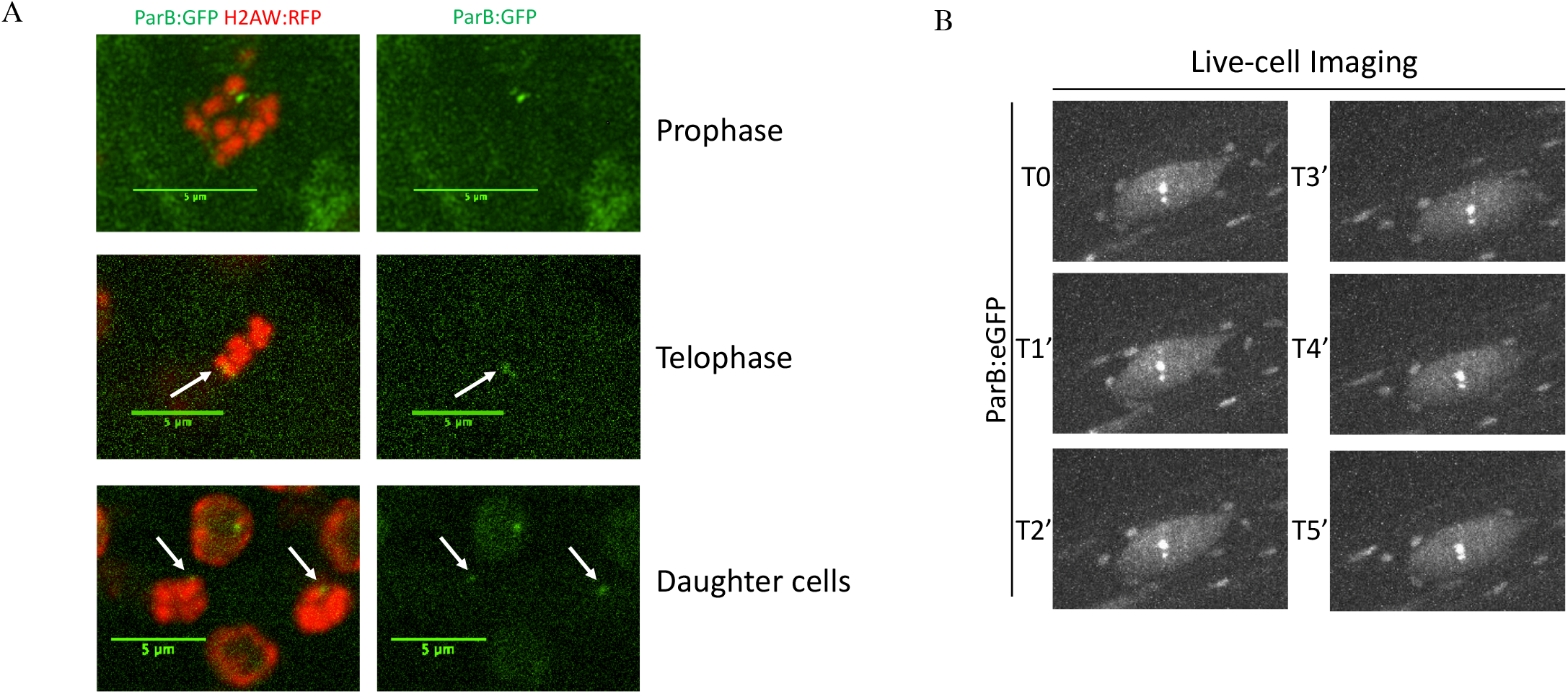
Monitoring pαrS-ParB:GFP in live during mitosis or during a time-course. A. Detection of *parS-*ParB:GFP foci (green) and Histone2A variant H2AW:RFP (red) in mitotic cells. B. ANCHOR system enables time-lapse tracking of a single-locus in live roots by confocal imaging. Time-lapse acquisition of *parS*-ParB:GFP signals (grey) in an endorepiicated root cell over 5 min.

Finally, we tested our ability to perform live-cell imaging of the *parS*-ParB:GFP signals *in planta.* We analysed *parS*-ParB:GFP signals in live using a Zeiss Cell Observer Spinning disk microscope (Figure 3B). Although bleaching could alter the signal detection in time, we were able to detect the ParB:eGFP signals during multiple time points and follow its relative nuclear position through time, as shown previously in human and yeast cells (Saad et al., 2014; Germier et al., 2017). Movies showing the *parS*-ParB:GFP signals detection in live meristematic or elongated cells can be find as supplementary data (Suppl. Movies 1 and 2). Altogether, our data demonstrates that the ANCHOR system is suitable for live-cell imaging *in planta.*

### Studying chromosome mobility using the ANCHOR system

It is now clear that higher-order organization of the chromatin exerts an important influence on genomic function during cell differentiation (Arai et al., 2017). For instance, in *Arabidopsis thaliana,* histone exchange dynamics was shown to decrease gradually as cells progressively differentiate (Rosa et al., 2013). However, how chromosomes and the chromatin fibre move during cell differentiation is not well studied in plants. We took advantage of our ANCHOR DNA labelling system to examine chromatin mobility during cell differentiation in Arabidopsis. In particular, we measured mobility of *parS*-ParB:eGFP foci in meristematic and differentiated cells from the root epidermis (Figure 5A) through live-cell imaging using confocal microscopy, and quantified the mobility using mean square displacement (MSD) analysis (Meschichi and Rosa, 2021). Interestingly, the chromatin mobility on meristematic cells was higher than in differentiated cells (Figure 5B, Suppl. Movies 1 and 2). These differences were statistically significant as shown by a much higher radius of constraint (Figure 5C). These results may support the idea that the chromatin in undifferentiated cells holds a more dynamic conformation (Meshorer et al., 2006; Rosa et al., 2013; Meschichi and Rosa, 2021). Additionally, this illustrates that the ANCHOR system can be used to monitor single-locus *in planta* and is suitable to unpick differences in chromatin mobility.

**Figure 5:**
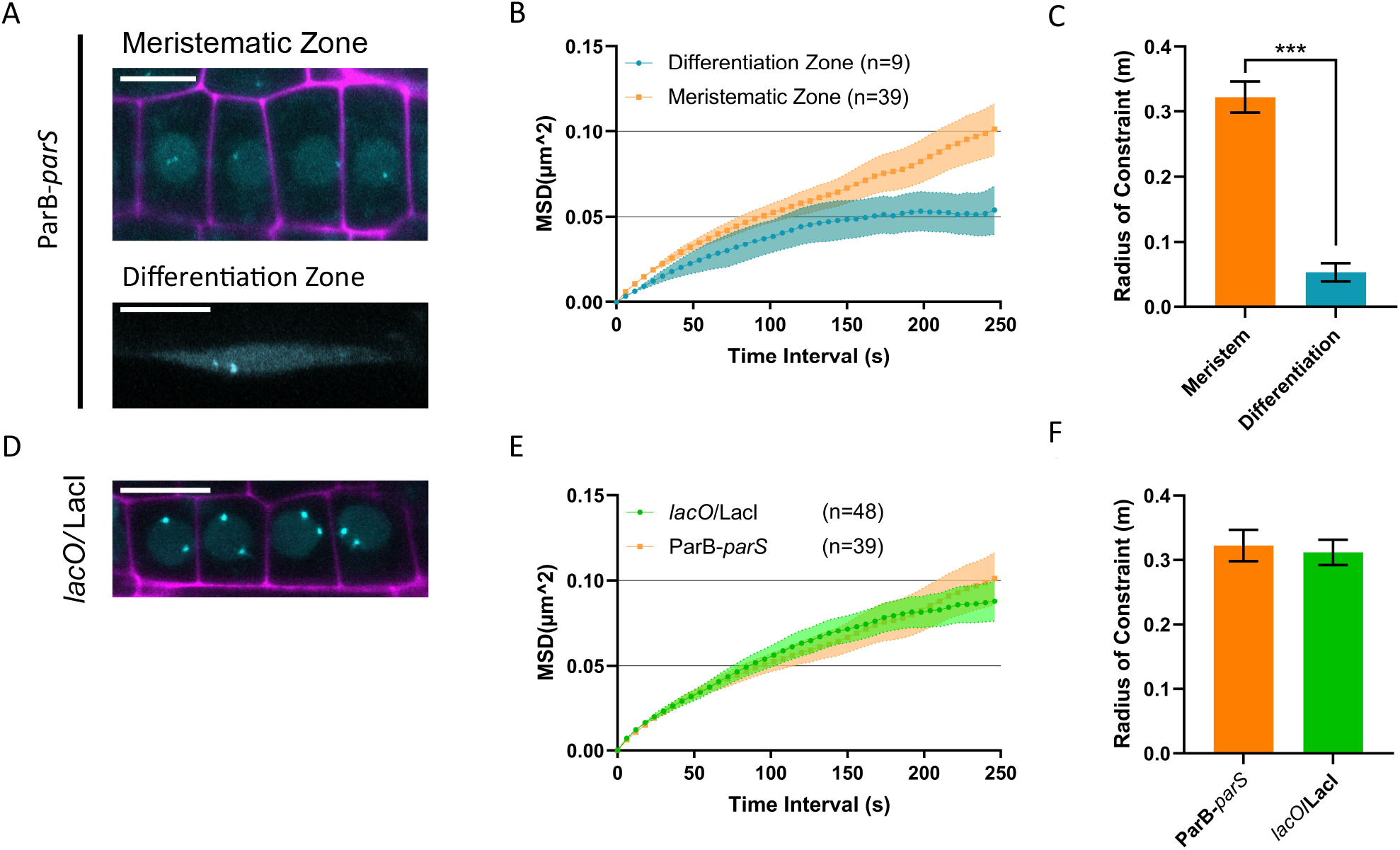
Analyzing chromatin mobility using the ANCHOR system. A. Representative images of ParB-*parS* line in meristematic (upper panel) and differentiation zone (bottom panel) showing nuclear signal with spots (cyan). Propidium Iodide (PI) staining (magenta). Bars = 10 μm. B. MSD analysis for ParB-*parS* lines based on time lapse experiments of nuclei in the meristematic and differentiated zone. 3D stacks were taken at бsec intervals for 5min. Values represent mean ± SEM from 54 and 9 cells, respectively. C. Calculated radius of constraint for MSD curves depicted in B. Values represent means ± SEM. Mann-Whitney U-test, ***P < 0.001. D. Representative image of/*lacO*/Lacl line in meristematic region showing nuclear signal with spots (cyan). Propidium Iodide (PI) staining (magenta). Bar = 10 μm. E. MSD analysis for *lacO*/LacI and ParB *-parS* lines based on time lapse experiment of nuclei in the meristematic zone. Values represent means ± SEM from 116 and 54 cells, respectively. F. Calculated radius of constraint for MSD curves depicted in E. Values represent means ± SEM.

Because until now single-locus dynamics in plants was mostly possible through the use of the *lacO* system (Figure 5D) we thought to compare chromatin mobility in meristematic cells using the ANCHOR and the *lacO* systems. Interestingly, both methods revealed very similar MSD curves with a comparable radius of constraint (Figure 5E,F) further validating the use of the ANCHOR system for studying chromatin mobility in plants.

## DISCUSSION AND PERSPECTIVES

In this manuscript, we describe a novel method to monitor a single-copy locus i*n planta.* Live cell imaging of genomic loci in plants was already possible through the use of different strategies, including zinc-finger, TALE, CRISPR/Cas9 based imaging or *lacO*/LacI. Unfortunately, these techniques have been so far restricted to follow the dynamics of moderately to highly repeated regions, or require the insertion of repeats, transforming the transgene into the target of choice for silencing (Watanabe et al., 2005; Hirakawa et al., 2015; Meschichi and Rosa, 2021; Rosa et al., 2013). In contrast, the *parS* sequence is only 1 kb long and could potentially be shorten to 200 bp (NeoVirtech, personal communication). Importantly, the ANCHOR system is free of repetitive sequences and several reports in yeast and animal cells have already demonstrated its innocuity to endogenous processes such as transcription and replication (Germier et al., 2018). This particularity makes the ANCHOR system very suitable to monitor single-copy genes in its native genomic environment. Indeed, recent approaches using CRISPR-Cas9 technology generated knock-in Arabidopsis plants (Wolter et al., 2018; Merker et al., 2020). Combining *parS* targeted insertion and having ParB:GFP being expressed using an independent transgene should allow robust single gene locus monitoring.

Another advantage of the ANCHOR approach is the possibility to use different combination of *parS*-ParB in the same time. ParB binding on *parS* sequence is indeed species-specific and several combination have successfully being used separately or simultaneously so far. In this study, we used a specific *parS*-parB, but additional specific combination could be used. In theory, up to three combinations could be used simultaneously (Saad et al., 2014, NeoVirTech peronnal communication), although an important preliminary work would be required for plant material preparation. For instance, two alleles from the same gene could be differentially labelled to monitor their potential associations while being expressed or silenced. This is an important question since previous observations suggest that allele aggregation could be part of the silencing mechanism (Rosa et al., 2013). These color combinations could also be used to follow the distance of two proximal regions during repair for example, as already shown in yeast (Saad et al., 2014) or to label borders of a genomic regions that can undergo different chromatin states during stress or development. This system will provide a useful tool to study the spatial organization and the dynamic behavior of chromatin at the single locus level.

### Competing interest statement

FG is an employee and FG and KB are shareholder of NeoVirTech. NeoVirTech did not have any scientific or financial contribution to this study. No other conflict of interest to declare. ANCHOR system is the property of NeoVirTech SAS, Toulouse, France. Any request of use should be addressed to

## Acknowledgments

We thank Guillaume Moissiard, Christophe Tatout and Aline Probst for fruitful discussions. We thank Chritel Llauro for her help with Nanopore sequencing. We also thank the flow cytometry facility, the microscopic facility, and the sequencing facility of platform BioEnvironnement of Perpignan University Via Domitia (Perpignan, France), as well as the Clermont-Ferrand Imagerie Confocale platform (Clermont-Ferrand, France). We acknowledge the imaging facility MRI, member of the national infrastructure France-BioImaging infrastructure supported by the French National Research Agency (ANR-10-INBS-04, «Investments for the future») FP was supported by the Agence Nationale de la Recherche (ANR), JCJC NucleoReg (ANR-15-CE12-0013-01) and by the French Laboratory of Excellence project TULIP (ANR-10-LABX-41 and ANR-11-IDEX-0002-02). MI was supported by the French National Research Agency (ANR-15-CE12-0012). M.M. is a member of the European Training Network “EpiDiverse” that receives funding from the European Union Horizon 2020 program under Marie Skłodowska-Curie grant agreement No. 764965. AM and SR were supported by the Swedish Research Council (Vetenskapsrådet) grant number 2018-04101. A.M., M.I., C.T., N.P., S.R. and F.P. are part of the European Cooperation in Science and Technology COST ACTION CA16212 INDEPTH.

## Author contributions

M.I., F.P. and S.R. designed the experiments. A.M., M.I., C.P. and F.P. performed the experiments. A.M., M.I., N.P., S.R. and F.P. analyzed the data. S.D., F.G., K.B. and M.M. participated in material preparation or analysing tools. F.P. wrote the paper and S.R. edited the paper. F.P. acquired main funding.

## Supplemental figures legend

**Figure S1: Chromatin states flanking the insertion site in T2F**

Snapshot of the chromatin states enriched in the region flanking the transgene insertion site in the line T2F (http://systemsbiology.cau.edu.cn/chromstates/UCSC.php).

**Figure S2: Pollen and trichome cell.**

Confocal images of the parS-ParB:GFP signal in a trichome cell (top panels) or in pollen cells (bottom panels). The images on the right are saturated to show the structures on the trichome or the envelope of the pollen.

**Figure S3: Col-localization of *parS*-ParB foci with H3Ac and H3K27me3 marks**

Detection of parS-ParB:GFP foci (Green) and post-transitionally modified histones (red) in fixed and isolated nuclei *from A. thaliana* Col-0 plants T2F. Nuclear DNA is labeled with DAPI (grey). Trimethylated H3K27 signals are shown in the panel A, while acetylated H3 are shown in the panel B. C and D panels show the relative intensity of each signals.

