## Supplementary figures and images for "ANCHOR, a technical approach to monitor single-copy locus localization *in planta*"

### Supplemental Figures

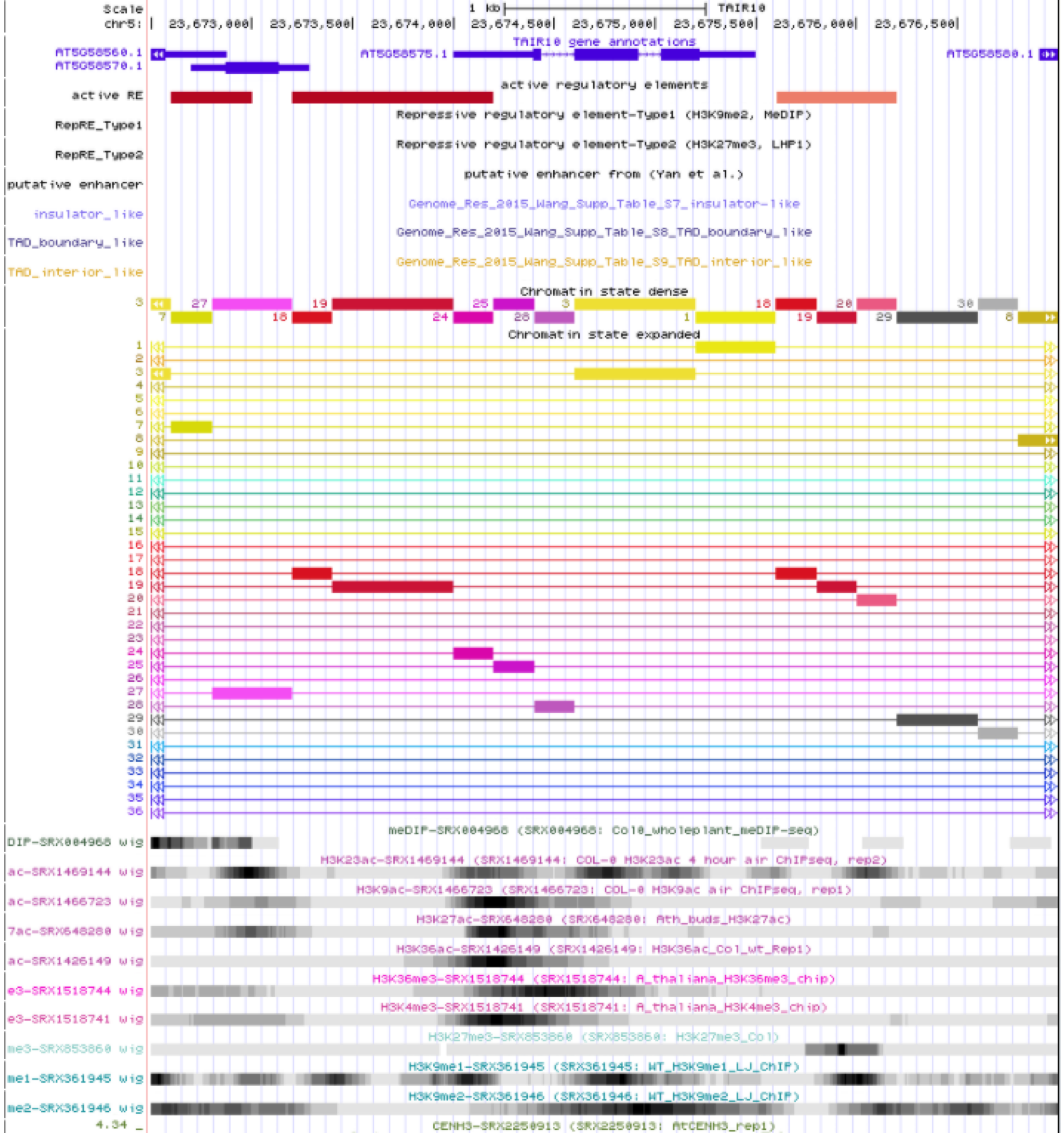

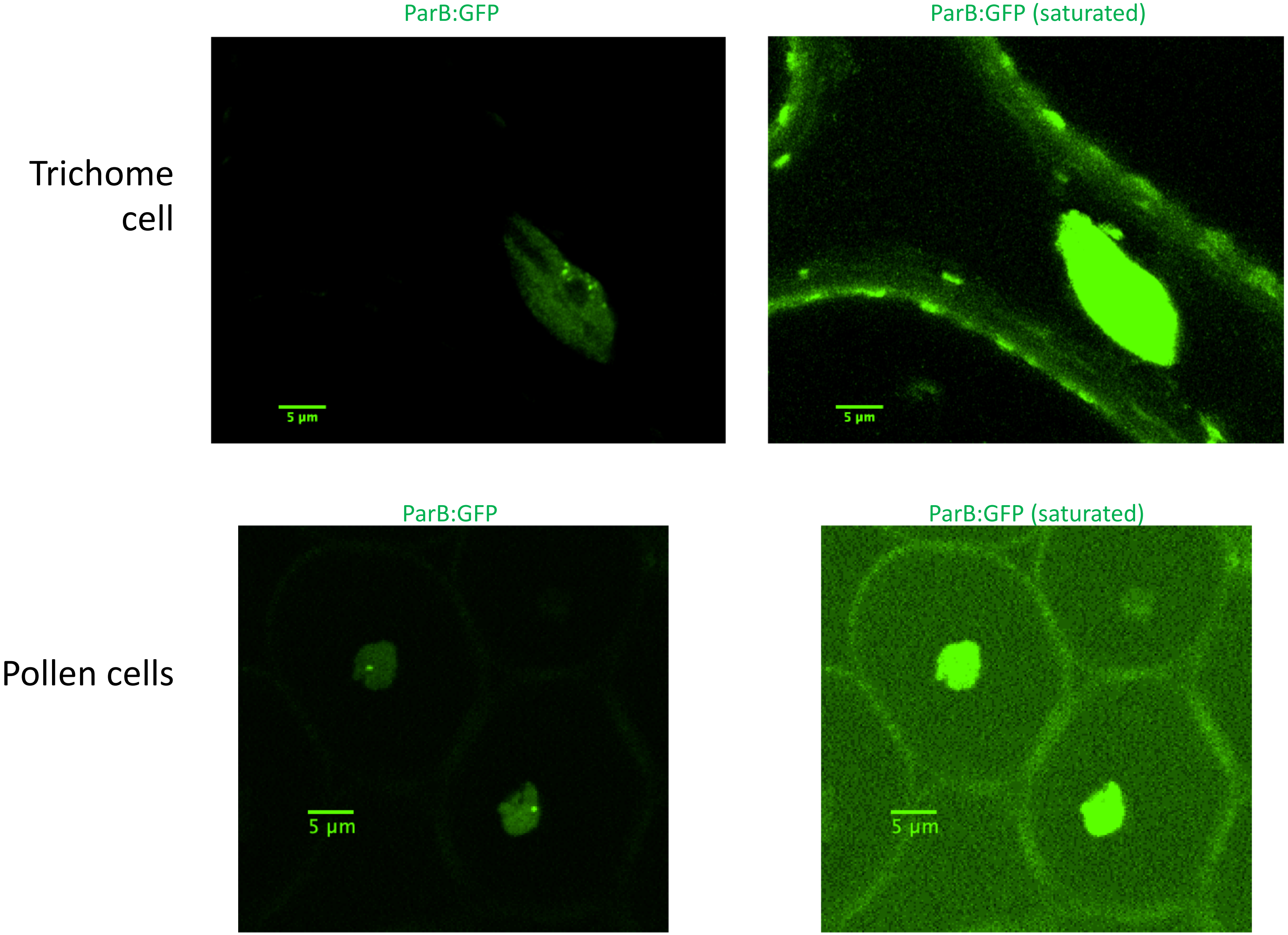

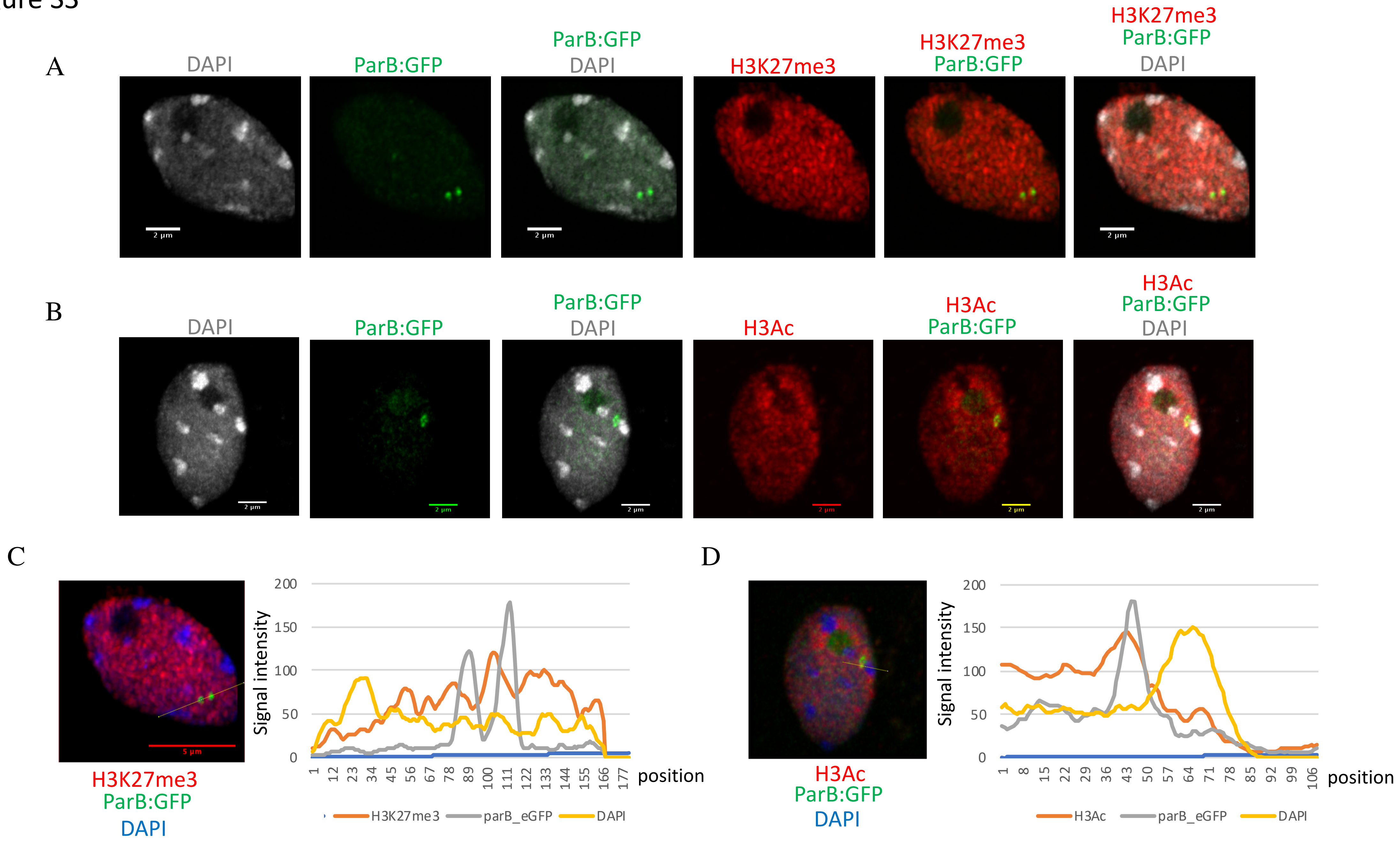
